# Humans and a subterranean rodent have contrasting biological rhythms

**DOI:** 10.1101/2023.10.05.561036

**Authors:** Otto Appenzeller, Clifford Qualls, Tim Bromage

## Abstract

Here we show that naked mole rats (NMR’s) have an extraordinary survival advantage. We base this statement on a spectral analysis of the time series of measured intervals in teeth of 3 species (NMR’s, Killer Whale, and Modern Humans). We used Fourier decomposition to analyze the variability of these intervals. We find these animals go through their long life without any of the age-associated diseases seen in humans such as osteoporosis or cancer and they show no signs of muscle atrophy or slowing of mobility. Global warming will delay its effects on these animals whereas for humans it is a clear and present danger.

## Introduction

Biological rhythm or circadian rhythm are bodily functions regulated by the internal clock. They control the cycles of repetitive events such as sleep and wakefulness, body temperature, hormone secretion, and many more.

Growth lines in archived biological materials such as teeth, also known as circadian rhythms, give evidence of metabolic health, and physiological function.[1]

Naked mole rats (Hereocephalus glaber) and humans are mammals. But, naked mole rats uniquely thermoregulate by changing the level of their burrows, whereas humans use shivering and other mechanism to maintain steady temperature.[2]

Here we report a comparison of growth lines on teeth of naked mole rats with growth lines on teeth of humans, and other mammals to determine how the statistical differences in spectra might explain their successful survival.

Unlike humans, naked Mole rats (NMR’s) cannot maintain a steady body temperature they huddle in warmer tunnels near the surface when cold; they do not shiver. They obtain their water through food; they do not drink. These animals are the longest living rodents; they live for approximately 30 years. Whereas mice (somewhat smaller) live only about 2 years. In a naked mole rat, the teeth can move independently.

## Material and Methods

See Figure 1 and Figure 2. In this study no animals were used.

**Figure 1.**
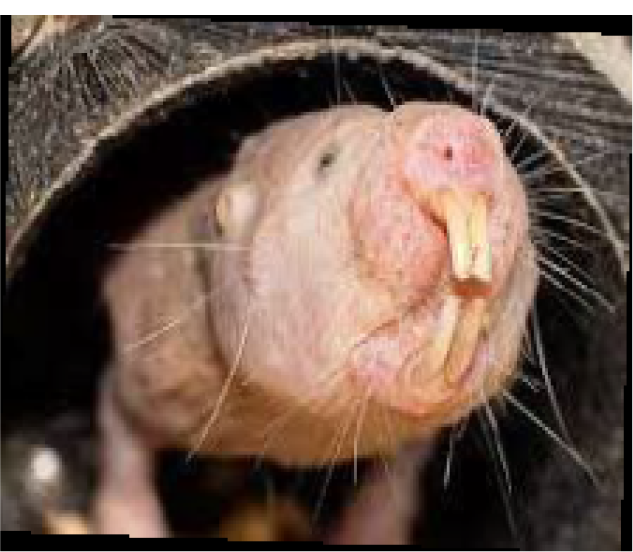
Left, naked mole rat. Right tooth of naked mole rat. The image on the right shows the growth lines of a naked mole rat tooth obtained scanning electron microscope (dark streaks).

**Fig. 2.**
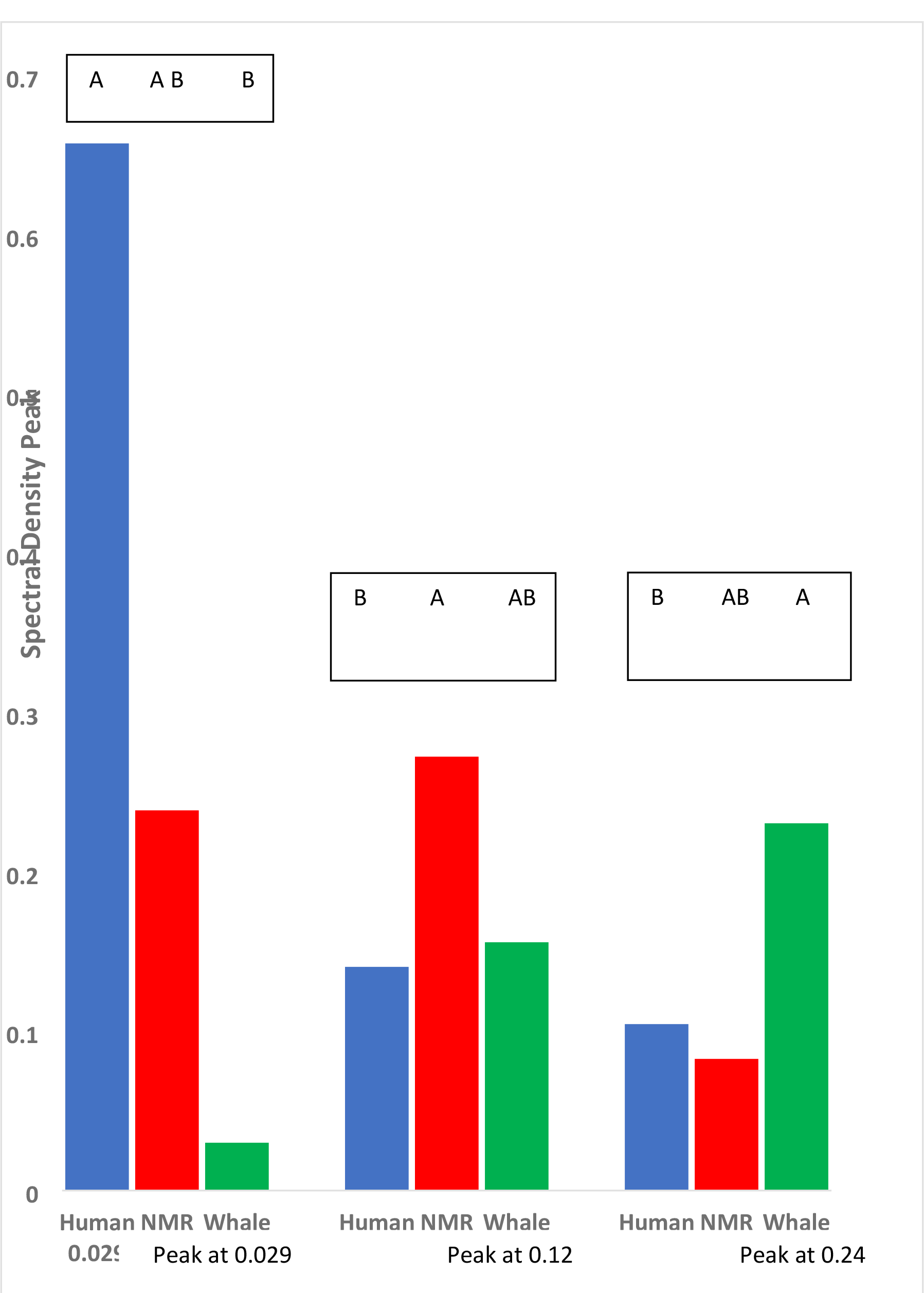
Spectral density peaks at given frequencies. Peaks observed at frequencies 0.029, 0.12 and 0.24.

Naked Mole Rat - NMR, and Whale at each given frequency are denoted by the letters A, B, C. The letter A marks the species with the largest peak at the given frequency. At a given frequency, those peaks of species marked by different letters are (statistically) significantly different (α=0.05) Spectral density values of species marked by AB are not significantly different than those with either A or B.

### Statistical Methods

Spectral analysis of the time series of measured intervals in teeth of 3 species (NMR, Killer Whale, Modern Human) includes the following. The Fourier decomposition the variability of these intervals gives the periodogram, the power at each frequency

[1,4]. The integration (a convolution with a spectral window) of the periodogram over frequencies λ gives estimates of the spectral density function *f*(λ) at each frequency λ. The spectral density estimates 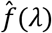obtained from these intervals in teeth for each species plotted versus the frequencies λ forms that species’ spectrum; these plots are overlaid in Fig. 1 for visual inspection. Statistical comparison among 3 species will be made using both low frequency/high frequency (LFHF) ratios as well as the locations of obvious spectral peaks in the 3 spectra.

### Comparison of LFHF ratios among the 3 species

The distribution theory of LFHF and of accumulated power in frequency bands is given in [2]. Briefly, the estimate of *f*(λ) is a smoothed and locally weighted average of the periodogram *I*(λ) :

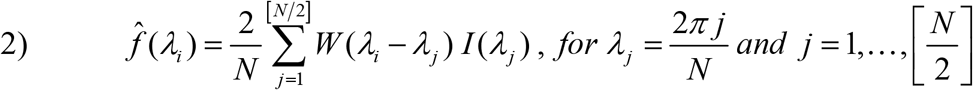

The symbol [*x*] represents the integer part of *x* and N= number of intervals, the length of the time series. Spectral density *f*(λ) is symmetric about λ = 0 by definition (definition not shown) and *I*(λ) is symmetric. Since *W*, called the spectral window, is taken to be symmetric, the estimated spectral density is symmetric, which allows one to plot the spectral density only for the nonnegative frequencies 0 *≤*λ*j ≤* π. Note that 4π/*N* = 2Δλ*j*, where the extra 2 represents the sum over the negative λ and the *y*-axis should also be scaled by dividing by 2π, and finally that (4π/*N*) ÷ 2π = 2/*N* is the coefficient of formula for *f*(λ). The power in any frequency band is estimated by the area under the curve (AUC):

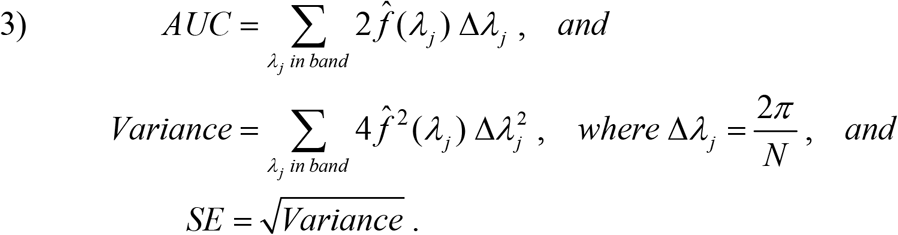

The distribution of the AUC estimator is based on the distribution of each estimated spectral densities *f*(λ*j*), which in turn depends on the effective degrees of freedom(EDF) of spectral window *W*(λ); see Koopmans [3, Table 8.1, page 279]. A typical approximation to the distribution of the estimator AUC given in eq. 3) above is a constant multiple of a Chi-square random variable:

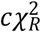where 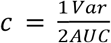and 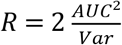with AUC and Var as computed in 3) above. The ratio 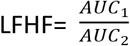 involves the ratio of independent Chi-squared random variable and so has a distribution of a constant multiple of a F-distributed random variable: 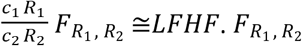. These distributional facts are enough to compute confidence intervals for AUC and P-values for comparisons.

### Comparison of the locations of spectral peaks among the 3 species

The comparison of locations (frequencies) of obvious spectral peaks in our spectra for our 3 species will be based on the power in narrow frequency bands isolating the identified frequency of a potential spectra peak; the use of the same band in all 3 species simplifies the comparison of the AUCs for that band. Since there are three species, there are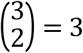 pairwise comparisons [4]. Typical of pairwise comparisons, our strategy at each identified frequency location (frequency band) will be to compare the species with the largest peak (and largest power) to the power of the other two species by computing the lower 95% confidence limit (LCL) and if either of the powers for the other two species are less than this LCL then the peak for such a species is (statistically) significantly less than the largest peak. For the third comparison, we compare the species with the second largest peak by computing its LCL to the species with the smallest peak. It is customary to label the largest peak and all of the peaks not statistically different than it with the letter ‘A’. The first peak different than the largest peak is labeled by ‘B’. If the first (largest) two peaks are different and the third peak is also different than the second peak then the third peak is labeled by ‘C’. The above strategy determines the pairwise differences, and computation of tail probabilities of the Chi-square distribution allows p-values to be given for each of the pairwise comparisons.

A difficulty with these pairwise comparisons is the wide disparity in sample sizes of the interval time series (Modern Human n=441; NMR n=59; Killer Whale n=11) which means the confidence intervals and one-sample test of hypotheses are much more accurate for Modern Man. Thus, the following modification of the pairwise comparison strategy is better: we order the species by sample size to determine the order of the pairwise comparisons.

## Discussion

Here we compare humans and naked mole rats to elucidate what makes these animals so extraordinary survivors.

[1]Naked mole rats (Heterocephalus glaber) are the longest-living rodents known, with a maximum lifespan of 30 years. Whereas a mouse, only slightly smaller, lives at most 4 years. They are not naked; they have some hair between their toes and on their ears. They are blind but they have eyes and have some sight left. They live underground and therefore lack the daily rhythms the sun provides for their bodily cycles such as feeding and sleeping. In this animal all clock genes peak at the same time, in the morning, whereas in all other terrestrial animals, they cycle every twenty-four hours.[2]

These animals go through their long life without any of the age-associated diseases seen in humans such as osteoporosis or cancer and they show no signs of muscle atrophy or slowing of mobility.[3]

Although they keep a steady body temperature, they cannot actively thermoregulate they dig deeper or shallower tunnels where they huddle to keep comfortable. They have up to 21 pups per litter; the female is the “head of the tribe”. Their fecundity far exceeds that of humans.[4]

Global warming will delay its effects on these animals whereas for humans it is a clear and present danger.

## Acknowledgements

We thank the NMHEMC Research Foundation for support and the NYU TEM access

